# Virus-derived Synthetic Riboswitches for Real-time Coronaviruses Sensing in Live Cells

**DOI:** 10.1101/2024.02.27.582433

**Authors:** Leiping Zeng, Lei S. Qi

## Abstract

Real-time sensing of viral infection in live cells are crucial for virology research and antiviral development. However, existing methods face challenges of low signal sensitivity and the necessity for viral manipulation and cell fixation. Here, we develop a <u>v</u>iral <u>ribo</u>switch (vRibo) approach that employs the viral replicase to induce transgene expression upon viral infection. The vRibo is designed to detect viral real-time transcription and replication in live cells, which in response triggers the translation of reporter and therapeutic genes. By integrating a viral packaging sequence, vRibo can be transmitted to neighboring cells through progeny virions, effectively acting as a ‘Trojan Horse’. The negative-strand vRibo elements demonstrated effective detection of several coronaviruses, including 229E and OC43, due to the conservation of cis-acting RNA structures across coronaviruses. Notably, vRibo functions as a dual-purpose system, acting both as an infection detector and inducible antiviral system. vRibo has the potential for basic virology research applications and can be adopted in improving the inducible expression of mRNA medicines for future coronaviruses.

## Introduction

Coronaviruses, a diverse family of single-stranded positive RNA viruses, include strains responsible for the common cold, such as 229E and OC43, and severe acute respiratory syndrome coronavirus 2 (SARS-CoV-2), the cause of the COVID-19 pandemic^1^. Although the World Health Organization (WHO) has declared the end of the COVID-19 global health emergency, the potential for newly emerging strains and rapid viral evolution remains a substantial concern. Therefore, continued monitoring and therapeutic development are essential to mitigate the threat posed by evolving coronaviruses.

Synthetic biology approaches that can sense and respond to virus infection and replication have high value for research, diagnostics, and therapeutics. They potentially offer precise identification and targeting of infected cells, improving our understanding of virus biology, and aiding the development of antiviral therapeutics. However, commonly used virus detection methods, such as quantitative polymerase chain reaction (qPCR) and antibody staining, require the lysis or fixation of cells and tissues. While some RNA sensors, including the eukaryotic toehold-based RNA sensor^2^, FlipGFP^3^, and ADAR-based RNA sensor^4^, have shown promise for viral detection in live cells, they often exhibit high background signals, low sensitivity, and are limited to certain applications.

We aimed to develop real-time RNA-based sensors that can detect a broad range of coronavirus strains in living cells. Compared to their protein or DNA counterparts, RNA-based sensors are less immunogenic and can be modularly combined with RNA vaccines or medicines, thus improving diagnostics and treatment. To identify suitable RNA-based sensors, we drew inspiration from the transcription mechanism of coronavirus genomic RNA (gRNA) and subgenomic mRNAs (sgmRNAs).

Coronaviruses use the replication-and-transcription complex (RTC) to specifically recognize gRNAs for replication and sgmRNAs for transcription. Notably, while replication of the coronavirus genome necessitates continuous RNA synthesis, transcription is a discontinuous process^5^. During transcription of the nested set of negative-strand subgenomic RNAs (sgRNAs), virus RTC performs a template switch at the site of transcription-regulating sequence (TRS) to add a common 5’ leader sequence (**Fig. 1a**). These sgRNAs subsequently serve as templates for the synthesis of multiple copies of sgmRNAs. Importantly, this process supposedly protects viral mRNAs from viral nonstructural protein 1 (Nsp1)-induced endonucleolytic cleavage and translation suppression of host mRNAs, providing a strategy for the efficient accumulation of viral mRNAs and viral proteins during infection^6,7^. Interestingly, defective interfering virus particles arise with reduced replication capability, which often contain random deletions but ubiquitously encode the 5’ and 3’ UTR structures to maintain the replication and packaging capabilities by the RTC^8–10^.

**Figure 1.**
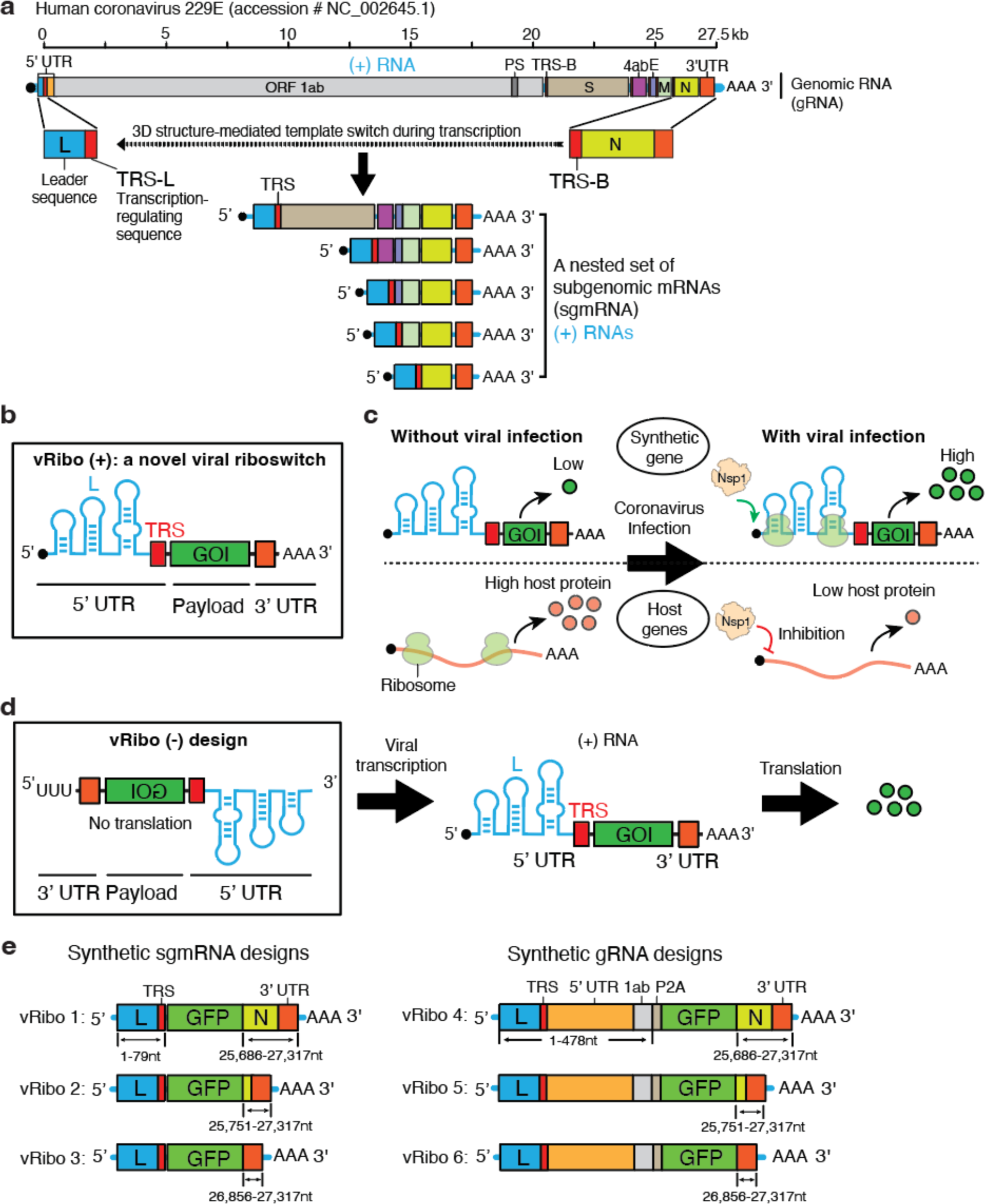
The design of coronavirus riboswitches. **(a)** A scheme depicting the structure of genomic and subgenomic RNAs of 229E and the viral RNA’s discontinuous synthesis strategy. **(b)** Design of the (+) viral riboswitch (vRibo +). **(c)** Proposed activities of vRibo-regulated synthetic genes and host genes, with and without viral infection. **(d)** Design of the (−) viral riboswitch (vRibo −). **(e)** Design of the viral riboswitches by mimicking the viral genomic and subgenomic RNAs.

Building on these insights, we developed a viral riboswitch (vRibo) that detects coronavirus infection in real time and mediates infection-inducible gene expression. Our vRibo was engineered by harnessing and linking the 5’ and 3’ UTR sequences of coronavirus RNAs to a transgene, such as a reporter GFP gene or a therapeutic gene. We demonstrated that the vRibo RNA can be transcribed and replicated by viral RTC and the transgene expression can be triggered by virus infection. When integrated with a packaging sequence, vRibo can be packaged into progeny virions and transmitted to uninfected cells. We further discovered vRibo can act on a broad range of coronaviruses and demonstrate its potential as a simultaneous virus infection detector by encoding GFP and an antiviral agent by encoding interferon (IFN). The vRibo will find uses for basic virology research and real-time viral detection as well as for improving the specificity of mRNA vaccines in the long term.

## Results

### Design of a Riboswitch for Coronavirus Transcription Detection

Coronaviruses synthesize genomic RNA (gRNA) and subgenomic mRNAs (sgmRNAs) in a nested, discontinuous manner^5^. Nonstructural protein 1 (Nsp1) regulates the translation of viral versus host mRNA. Nsp1 induces endonucleolytic cleavage of host capped mRNAs and inhibits translation by binding to the ribosomal 40S subunit, blocking the mRNA entry tunnel^6,7^. However, it has been demonstrated that the leader sequence (L) of the 5’ UTR prevents Nsp1 from suppressing viral mRNA translation^7^. We hypothesized that this could be harnessed as a riboswitch mechanism for conditional expression of desired reporter and therapeutic genes, dependent on the infection status of the cell (**Fig. 1b-c**).

While the composition of the 5’ UTR, including the leader sequence and transcription-regulating sequence (TRS), is known, the essential sequence of the 3’ UTR required for effective transcription and translation remains uncharacterized^11^. To design riboswitches capable of detecting viral infection, we utilized regulatory elements from the human coronavirus 229E. We developed six virus riboswitch designs (vRibo 1-6), with three mimicking the shortest viral sgmRNA and the others emulating the viral gRNA (**Fig. 1e**). All designs encoded a GFP reporter, but with different flanking 5’ and 3’ UTR sequences. We proposed that negative RNA versions of the designs may offer a more stringent expression system, as they require transcription by the RTC first, significantly reducing baseline translation (**Fig. 1d**).

To test the effectiveness of different designs, we transfected HEK293T cells stably expressing the human APN protein (293T/hAPN) with each plasmid DNA encoding a different vRibo reporter and subsequently infected cells with 229E coronavirus. All designs were expressed under a CMV promoter with an incorporated HDV ribozyme tail to facilitate accurate RNA processing at the 3’ end (**Supplementary Fig. 1a**). While none of the positive designs showed virus infection-induced reporter expression (**Fig. 2a**), all negative designs demonstrated a strong induction (**Fig. 2c**). Riboswitch designs with the shortest 3’ UTR (3 and 6) displayed better activation upon viral infection.

**Figure 2.**
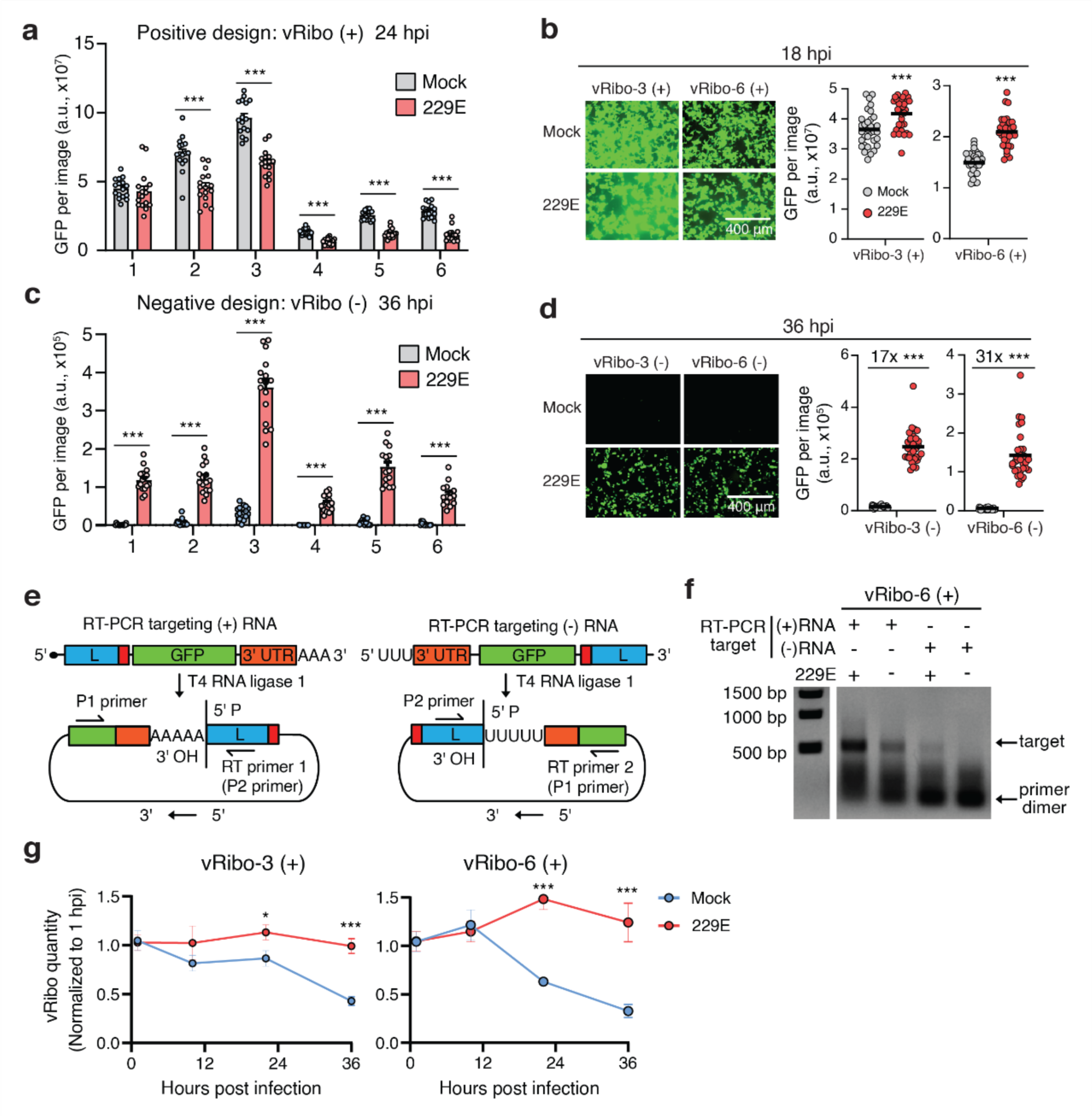
vRibos can be transcribed and replicated by coronavirus infection. **(a-d)** The expression of GFP reporter from vRibo (+) **(a-b)** and vRibo (−) **(c-d)** with or without 229E infection. The GFP intensity is the integrated signal calculated using the IncuCyte Live-cell Imaging System. 4 independent biological replicates were performed (4 images were taken for **a** and **c**, and 8 images for **b** and **d** per biological replicate). The bar represents the average of the group, while each circle represents an individual image. **(e)** The method for detection of (+) and (−) RNA transcript using RT-PCR. **(f)** The gel image of the RT-PCR products for detection of the (−) RNA transcribed from (+) vRibo after infection at 24 hpi. **(g)** The replication of vRibo-3 (+) and vRibo-6 (+) by virus infection. The 293T/hAPN cells were transfected with the in vitro transcribed RNAs and infected with 229E. Total cellular RNA was collected at 1, 10, 22, and 36 hpi. The vRibo RNA was quantified by RT-qPCR and normalized to 1 hpi. 3 independent biological replicates were performed with 3 technical repeats per biological replicate. Data are presented as mean ± s.e.m. P values were calculated by two-tailed Student’s t tests. n.s., not significant; *P < 0.05; ***P < 0.001.

We further characterized the GFP expression dynamics for vRibo-3 and vRibo-6 for both positive and negative designs and confirmed that the negative designs exhibited 17- and 31-fold of reporter expression, respectively, upon 229E infection (**Fig. 2b, 2d**, **Supplementary Fig. 1b-c**).

### Characterization of vRibo for Coronavirus Transcription Detection

To confirm this finding, we transfected cells with in vitro transcribed (IVT) positive vRibo. Following 229E infection, the cells displayed significantly higher reporter expression than the mock-infected cells (**Supplementary Fig. 1d**). While the negative vRibo needs to be transcribed into positive strand for protein translation (**Fig. 1c**), 229E infection induces the reporter expression from the negative vRibo, implying it is transcribed into positive sense RNAs by the viral RTC (**Fig. 2d, Supplementary Fig. 1e**).

T7 RNA polymerase produces ∼1% of the reverse complementary strand of the IVT RNAs^12^, to determine whether the RNA sensors are transcribed following viral infection, we followed a previously established protocol^13^ (**Fig. 2e**). Briefly, total cellular RNA was decapped and ligated head-to-tail by T4 RNA ligase 1. A primer specific for each RNA strand was used for reverse transcription through the head-to-tail sequence, which was then amplified using PCR. As hypothesized, cells transfected with positive vRibo RNA can be transcribed to negative-strand RNA following viral infection (**Fig. 2f**). Furthermore, we found that the virus infection can replicate the viral riboswitches, maintaining their abundance, while the IVT RNA decreased in mock-infected cells (**Fig. 2g**).

### vRibo Can Encode Bicistronic Genes and Package into Progeny Virions

The transcription regulation of coronaviruses allows them to synthesize sgmRNAs from gRNA, guided by the TRS sequence^5^. Building upon this understanding, we designed vRibos to encode a second protein that could be activated during viral infection (**Fig. 3a**). We incorporated a subgenomic RNA encoding mRuby3 into both the vRibo-3 and vRibo-6 designs, which mimic subgenomic and genomic RNA, respectively. Following transfection into HEK293T cells, GFP expression was observed. However, mRuby3 expression was only detected in cells infected by the 229E coronavirus (**Fig. 3b-d, Supplementary Fig. 2a-c**). We noted that the genomic RNA design (vRibo-6) expressed significantly higher levels of mRuby3 compared to the subgenomic RNA design (vRibo-3). Moreover, While GFP protein expression was reduced by infection compared to the mock condition at 48 hpi when vRibo encodes solely the GFP (Fig. S1b), its expression increased when encoding bicistronic proteins (GFP and mRuby) (**Fig. 3c**).

**Figure 3.**
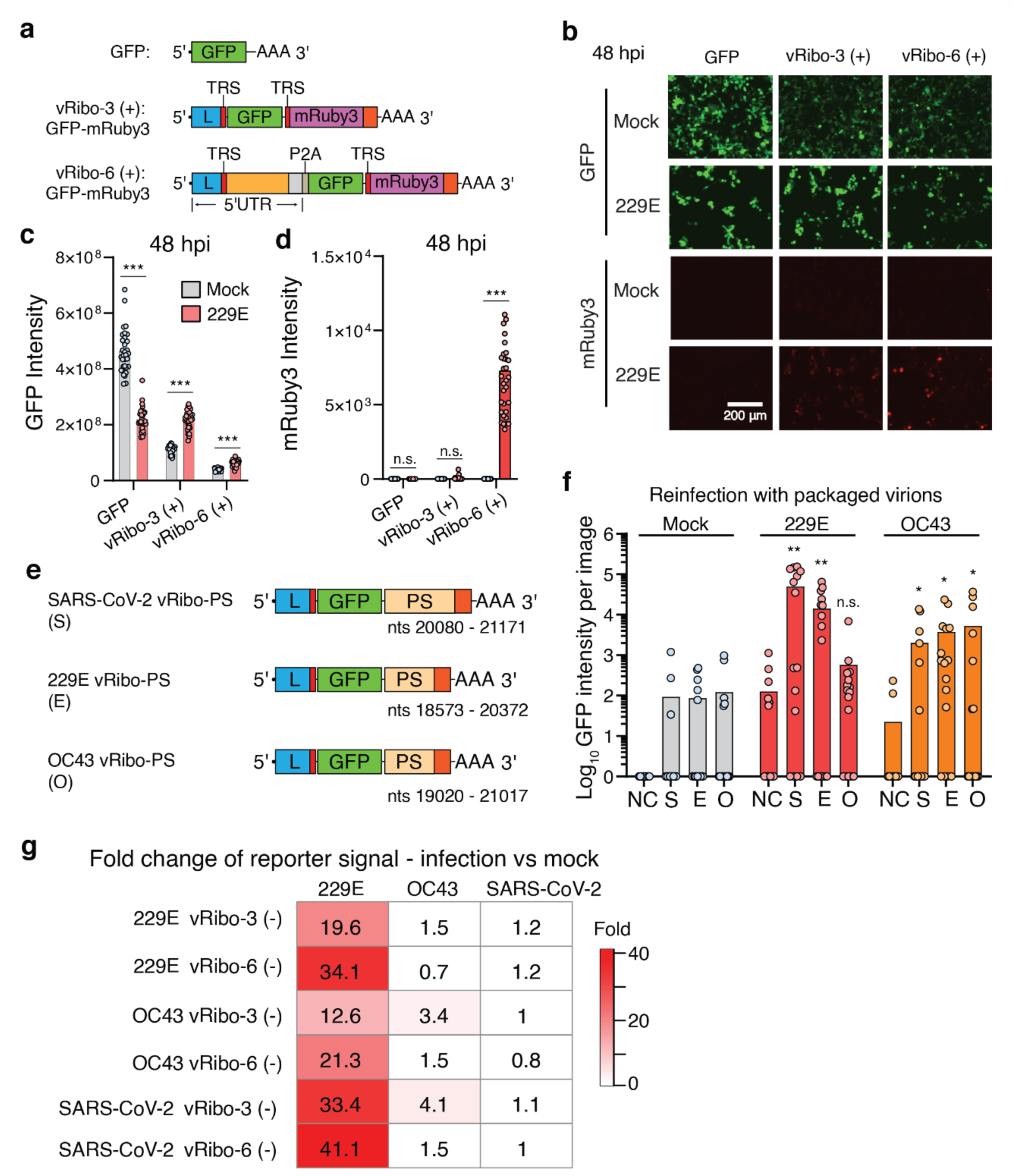
vRibos can encode additional proteins and be packaged into progeny virions for neighboring cell transmission. **(a)** The genomic and subgenomic vRibo designs encoding both GFP and mRuby3. **(b-d)** The expression of GFP and mRuby3 in the 293T/hAPN cells with or without 229E infection at 48 hpi. 4 independent biological replicates were performed with 8 images per biological replicate. The bar represents the average of the group, while each circle represents an individual technical repeat. P values were calculated by two-tailed Student’s t-tests. **(e)** The design of vRibos of SARS-CoV-2, 229E, and OC43 with a packaging sequence. **(f)** The GFP signal observed in 293T/hAPN cells upon re-infection with virions collected from the media of 293T/hAPN cells without transfection (negative control, NC) or transfected with vRibos (S, E, O) and either mock-infected or infected with 229E or OC43. 4 independent biological replicates were performed with 4 images per biological replicate. The bar represents the average of the group, while each circle represents an individual image. P values were calculated by one-tailed Student’s t-tests. **(g)** The fold activation of the GFP signal in the infected vs mock cells expressing vRibo (−) of 229E, OC43, and SARS-CoV-2, of which the source data is plot in Fig. S3e-g. n.s., not significant; *P < 0.05; **P < 0.01; ***P < 0.001.

We performed experiments to detect the synthesis of the subgenomic RNA from the vRibos using reverse transcription PCR (RT-PCR) (**Supplementary Fig. 2d**). A smaller secondary product was only generated by vRibo-6 in the virus-infected sample (**Supplementary Fig. 2e**). Sanger sequencing confirmed the association of the leader sequence with the mRuby3 coding sequence, as orchestrated by the viral RTC (**Supplementary Fig. 2f**)

Coronaviruses genome RNA can be packaged into progeny virions using a packaging sequence (PS)^14,15^. To explore this feature, we integrated a validated PS from SARS-CoV-2 into the vRibo-3 design and introduced potential PSs of 229E and OC43 into their corresponding vRibo-3 designs (**Fig. 3e**, **Supplementary Fig. 2g**). Upon virus infection with either 229E or OC43, we collected the media to infect fresh 293T/hAPN cells. The expression of the GFP reporter in the reinfected cells revealed that both SARS-CoV-2 and 229E vRibo-3 designs could be packaged by 229E virions. However, OC43 vRibo-3 was not packaged. Interestingly, all three riboswitches were successfully packaged by OC43 virions (**Fig. 3f**).

### vRibo Can Detect and Respond to Broad Coronaviruses

In addition to the 229E vRibos, we also designed 6 negative-strand vRibo RNAs each for OC43 and SARS-CoV-2, using the same engineering strategy (**Supplementary Fig. 3a, 3c**). These were tested in MRC-5 and 293T/hACE2 cells, respectively. Among the OC43 vRibos, those with the longest 3’end sequence, which includes the N protein-coding sequence and the 3’ UTR, displayed the most substantial fold of activation in response to virus infection (**Supplementary Fig. 3b**). However, the SARS-CoV-2 vRibos showed only mild induction of reporter expression due to their high baseline expression in uninfected cells (**Supplementary Fig. 3d**). Interestingly, we found these vRibos capable of cross-activation upon testing with infection of the three different coronaviruses (**Fig. 3g, Supplementary Fig. 3e-g**). The 229E infection successfully triggered reporter expression from the vRibos of all three viruses. In contrast, the OC43 infection only activated reporter expression from OC43 vRibo-3 and SARS-CoV-2 vRibo-3. Notably, infection with SARS-CoV-2 did not appear to activate reporter expression from any of the vRibo designs. The non-specific activation of vRibo by various viruses upon infection may suggest a certain degree of conservation in the cis-elements and the replication machinery of these coronaviruses.

### vRibo has Potential for Coronavirus Detection and Antiviral Responses

The vRibo system can hijack the virus replication machinery to enable self-transcription. When vRibo is expressed as a negative-strand RNA, it can be converted into a positive strand, thus activating transgene expression upon viral infection. By encoding a reporter gene, vRibo can be employed to detect virus infection. We found that 293T/hAPN cells expressing 229E vRibo-3 (−) could detect 229E virus infection at titers exceeding 800 TCID50 (**Fig. 4a**).

**Figure 4.**
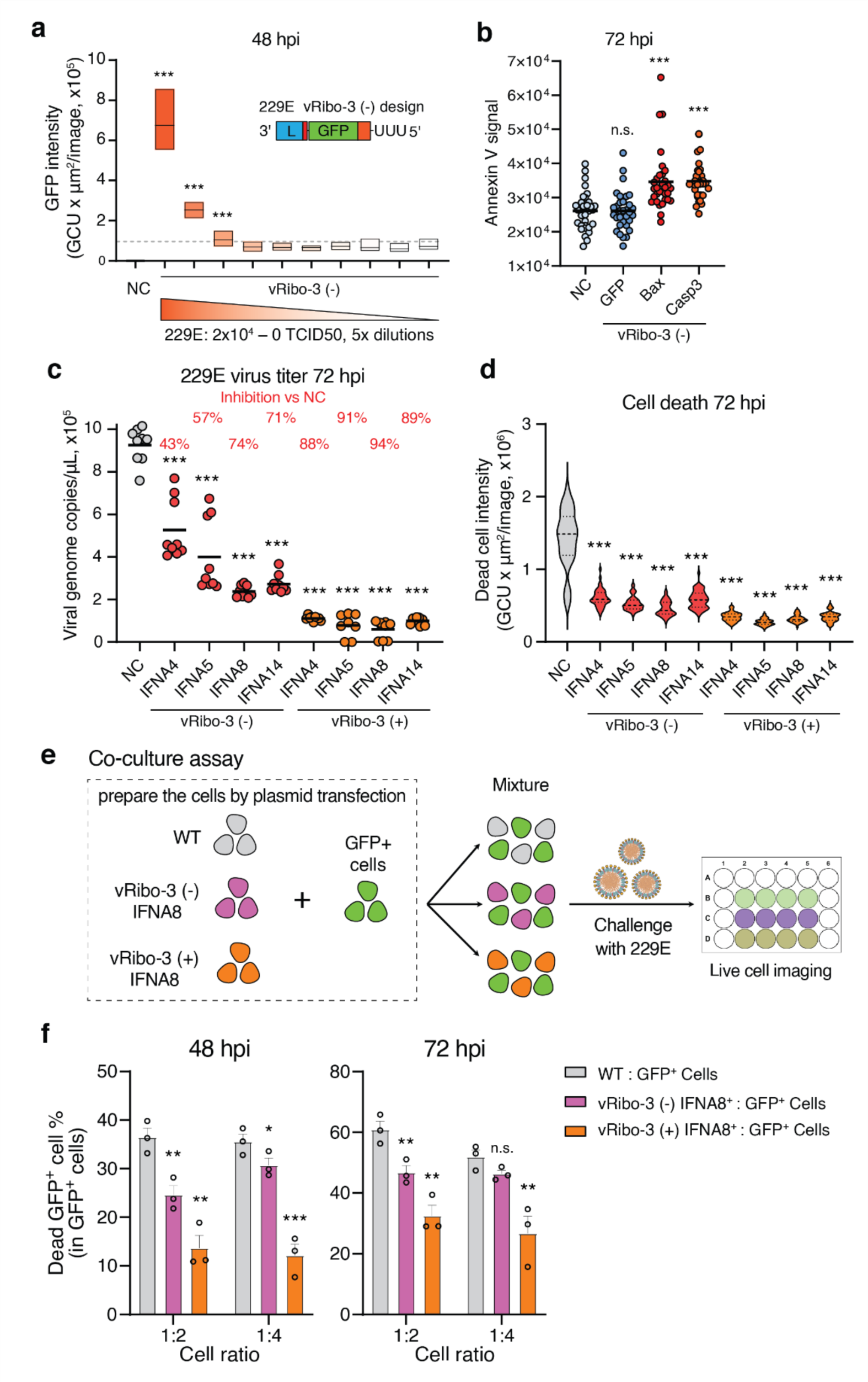
vRibos serve as live-cell virus detector and infection-activated antiviral strategy. **(a)** The GFP intensity of 293T/hAPN cells infected with or without 229E virus at various concentrations. Data are presented as box plots of each group with 4 independent biological replicates and 8 technical repeats per biological replicate. Box plots show the median (center line) with bounds of 25-75th percentiles. Data are presented as box plot showing the median (center line) with bounds from min to max. P values were calculated by two-tailed Student’s t-tests. **(b)** The apoptosis signal (Annexin V) of the cells that were transfected with or without an vRibo encoding an apoptosis inducer (Bax or Caspase3) and infected with 229E virus. 4 independent biological replicates were performed with 8 images per biological replicate. **(c)** The 229E virus titer in the media of the cells that were transfected with or without a vRibo expressing an interferon alpha subtype in either positive (+) or negative (−) strand and infected with the 229E virus. 3 independent biological replicates were performed with 3 technical repeats per biological replicate. The bar represents the average of the group, while each circle represents an individual technical repeat. P values were calculated by two-tailed Student’s t-tests. **(d)** The death signal (Cytotox green dye) of the cells. 3 independent biological replicates were performed with 16 images per biological replicate. Data are presented as violin plot. P values were calculated by two-tailed Student’s t-tests. **(e)** The co-culture assay scheme for testing the protective effects of vRibo-3 (+) or vRibo-3 (−) IFNA8 on neighboring cells. **(f)** The death rate of GFP^+^ cells in total GFP^+^ cells. 3 independent biological replicates were performed. The center line of each figure indicates the median. Data are presented as mean ± s.e.m. P values were calculated by one-tailed Student’s t-tests. NC, no transfection negative control; n.s., not significant; *P < 0.05; **P < 0.01; ***P < 0.001.

In addition to virus detection, vRibo could be used as a virus infection-activated antiviral system when encoding apoptosis inducers or interferons (IFN) (**Fig. 4b-d, Supplementary Fig. 4a-d**). When apoptosis inducers (Bax or Caspase 3) were encoded in the negative-strand vRibo-3, viral infection triggered transgene expression (**Supplementary Fig. 4b**) and markedly increased cell apoptosis, compared to untransfected cells or cells transfected with the GFP encoding vRibo-3 (**Fig. 4b**).

We encoded interferon alpha (IFNα) subtypes 4, 5, 8, and 14, known to effectively suppress SARS-CoV-2 replication^16^, in both positive and negative strands of the 229E vRibo-3 (**Supplementary Fig. 4c**). Compared to untransfected control cells, cells with positive-strand vRibo-3 RNAs effectively inhibited virus replication and reduced cell death as expected (**Fig. 4c-d**), with virus titers dropping by ∼ 80% and ∼ 90% at 48 and 72 hour-post-infection (hpi), respectively (**Fig. 4c, Supplementary Fig. 4d**). This underscores the efficacy of these IFNs against the 229E virus.

As expected, since transgene expression from the negative-strand vRibo-3 is dependent on virus infection, its onset is slower and less potent compared to positive-strand vRibo-3 RNAs. Among the four IFNα subtypes, negative-strand vRibo-3 encoding IFNA8 showed the most potent antiviral effect, reducing virus titers by 67% at 48 hpi and 74% at 72 hpi.

Interferons act as ‘alarm’ systems, alerting neighboring cells to prime their defense mechanisms against viral infections. Hence, vRibo-3 RNAs encoding IFNs may not only inhibit virus replication in the infected cells but also serve as a virus infection-activated alarm system for healthy neighboring cells. To evaluate this hypothesis, we designed a co-culture assay (**Fig. 4e**). We mixed wild-type, negative-strand vRibo-3, or positive-strand vRibo-3 IFNA8 expressing cells with GFP positive cells in a ratio of 1:2 or 1:4, and infected the cell mixtures with the 229E virus. Total cell death signal and GFP+ cell death rates were measured at 48 and 72 hpi. Both total and GFP+ cell death rates were highest when wild-type and GFP+ cells were co-cultured (**Fig. 4f, Supplementary Fig. 4e-f**). As expected, cell expressing positive-strand vRibo-3 IFN8 reduced overall cell death and effectively protected GFP+ cells from viral destruction. Similarly, cells expressing negative-strand vRibo-3 IFN8 also provided protection for GFP+ cells. These findings highlight the potential of our negative-stranded viral riboswitches as a novel responsive antiviral strategy, whereby transgene expression is triggered solely upon viral infection.

## Discussion

We’ve developed a virus riboswitch (vRibo) leveraging the RNA synthesis mechanism of coronaviruses to detect and respond to viral infection (**Supplementary Fig. 5**). The vRibo, incorporating viral 5’ UTR and 3’ UTR, can be transcribed and replicated upon virus infection, increasing transgene expression in both positive- and negative-strand formats. Moreover, integrating a TRS and an additional transgene allows the expression of a subgenomic RNA and an extra protein. By inserting a packaging sequence, the vRibo can be packaged into progeny virions and transmitted to other cells, thereby acting as a Trojan horse within the virus. Importantly, these vRibos exhibit broad activity across various coronaviruses.

The vRibo, when encoding a GFP reporter, effectively detects virus infection in live cells, showing potential for in vivo live-cell imaging of coronavirus infection and replication. A significant advantage of this system is that it circumvents the need to manipulate the large virus genome. Furthermore, it displays promising prospects for development into a virus-infection-activated antiviral strategy by using the negative strand-based vRibo, exhibiting protection against virus-induced cell death in both infected and uninfected cells. We anticipate that vRibos can be combined with RNA vaccines to further advance RNA therapeutics in infectious diseases, thus contributing to both virology research and the development of better antiviral therapeutics.

However, there are areas for improvement in the vRibo system, particularly in enhancing sensitivity for virus titer detection and minimizing background expression in mock-infected cells. We also anticipate hurdles with in vitro transcription of negative-stranded RNAs due to a long U stretch at the RNA molecules’ start. Overcoming these challenges could substantially enhance the vRibo’s efficacy.

Additionally, the concept of vRibo expands beyond coronaviruses, offering an exciting opportunity for the design of similar synthetic biology toolkits for other RNA viruses. By ingeniously exploiting viral replication or other viral machineries, we could develop sophisticated systems that can detect and respond to virus infections, advancing both virology studies and antiviral solutions. We believe this exploration with vRibos in coronaviruses offers a new and inspiring concept in the field synthetic virology, whereby extracting regulatory components from viruses and designing synthetic versions, we may create applications tailored for better virus detection and therapeutics.

## Methods

### Cell cultures, viruses, and plasmids

Human embryonic kidney (HEK293T, Cat# CRL-3216), MRC-5 (Cat# CCL-171), and Vero E6 (Cat# CRL-1586) cells were purchased from ATCC and cultured in 10% fetal bovine serum (Alstem # FB500) in DMEM (Gibco # 10569044). A HEK293T cell line stably expressing the human aminopeptidase N (APN) was made by lentivirus transduction and a monoclonal cell was selected for infection assays by human coronavirus 229E and OC43. The cell line was named 293T/hAPN. A HEK293T cell line stably expressing the human angiotensin-converting enzyme 2 (ACE2) was made using the same method and named 293T/hACE2. All cell lines were incubated at 37°C in a 5% CO_2_ atmosphere.

Human coronavirus 229E (Cat# VR-740, ATCC) and OC43 (Cat# VR-1558, ATCC) were amplified and tittered using MRC-5 cells as described previously^18^. All experiments involving the human coronavirus 229E and OC43 were performed in a Biosafety Level (BSL) 2 laboratory. All experiments involving the SARS-CoV-2 were performed in the BSL3 facility at Stanford University.

APN plasmid is purchased from Genscript (Cat#OHu16562C). ACE2 plasmid is from Addgene (Cat#1786). All the other plasmids used in this study are cloned by following the standard protocol of Infusion HD (Takara).

### Quantitative reverse transcription PCR (RT-qPCR)

As described previously^18^, RT-qPCR was performed with RNA standards synthesized using a TranscriptAid T7 High Yield Transcription Kit (Thermo Scientific Cat# K0441) from the DNA templates generated by RT-PCR from the SARS-CoV-2 and 229E viral RNA. For each sample, total RNA was isolated using a Quick-RNA Viral kit (Zymo Research Cat# R1035). RT-qPCR was performed using the GoTaq Probe 1-Step RT-qPCR System (Promega Cat# A6120). RT-qPCR primers and probes were listed in Supplementary Data 2.

### In vitro transcription (IVT)

The virus riboswitch (vRibo) RNAs were synthesized using a TranscriptAid T7 High Yield Transcription Kit (Thermo Scientific Cat# K0441) from the DNA templates generated by PCR. The template DNA was eliminated using DNase I. The positive sense RNAs were capped with vaccinal capping enzyme (NEB # M2080S) and methyltransferase (NEB# M0366S) by following the vendor’s manuscripts.

### vRibo activity test

293T/hAPN cells were seeded at 100 k per well in a 24-well plate one day in advance and transfected with the vRibo. After 3 hours, the cells were inoculated with 229E or OC43 at an MOI of 1.0. The cells were scanned for GFP expression using the IncuCyte live cell imaging system for 3 days. GFP intensity was calculated by using the IncuCyte Live-cell Imaging System.

### vRibo RNA transcript detection

As described previously^13^, the T7 RNA polymerase will generate 1% reverse transcript. To prove the vRibo RNA is transcribed by virus replicase, an RT-PCR method was performed as described previously^13^. The vRibos were transfected into 293T/hAPN cells and infected with 229E. The total RNA was isolated at 24 hpi and decapped with mRNA decapping enzyme (NEB# M0608S) and ligated head to tail by using T4 RNA ligase 1. Then RT-PCR with the indicated primers (P1 primer (i.e., RT primer 2): 5’-GGGATCACTCTCGGCATGG-3’; P2 primer (i.e., RT primer 1): 5’-AGACACAAAGTCTAAAAAGCAACT-3’) was used to testify whether the anti-sense strand of the vRibo RNA is generated.

### vRibo RNA replication detection

The vRibo RNA was prepared by IVT as described previously. 293T/hAPN cells were seeded one day in advance at 200 k per well in a 24-well plate and transfected with 500 ng of IVT RNA per well using lipofectamine MessegerMax (Thermo # LMRNA003). After 1 hour, the cells were infected with 229E at an MOI of 1.0 and were lysed with RNA lysis buffer (Zymo# R1035) at 1, 10, 22, and 36 hpi. Total RNA was isolated. The vRibo RNA was quantified using RT-qPCR with a primer pair (5’-ACGACGGCAACTACAAGACC-3’ and 5’-TTGTACTCCAGCTTGTGCCC-3’). RT-qPCR was performed using the iScript™ cDNA Synthesis Kit and iTaq Universal SYBR Green Supermix (Bio-Rad) and run on a Biorad CFX384 real-time system (C1000 Touch Thermal Cycler), according to the manufacturer’s instructions. The relative vRibo RNA abundance was normalized to the *GAPDH* internal control which was quantified with a primer pair (5’-GTCTCCTCTGACTTCAACAGCG-3’ and 5’-ACCACCCTGTTGCTGTAGCCAA-3’). The relative RNA abundance of each timepoint was normalized to the 1 hpi samples.

### Progeny virion package test

The 293T/hAPN cells were transfected with vRibo expressing plasmid and inoculated with 229E or OC43 at an MOI of 0.1. At 48 hpi, the supernatant was collected by spinning at 4000 g for 5 min and filtering through 0.45 μm and used to inoculate fresh 293T/hAPN cells. GFP expression was measured using the IncuCyte system for 2 days.

### Co-culture assay

The GFP^+^ cells were prepared by transfecting the 293T/hAPN cells with a GFP-expressing construct. The 293T/hAPN cells were not transfected or transfected with positive or negative strand vRibo encoding IFNA8 and mixed with the GFP^+^ cells at a ratio of 1:2 or 1:4. The cells were challenged with 229E at an MOI of 0.1. The Cytotox red dye (Sartorius # 4632) for counting dead cells was added to the media at a final concentration of 100 nM. The dead cells were counted using the IncuCyte system for 3 days.

### Data analysis

Data visualizations (graphs) were performed in GraphPad Prism software version 9. P values were calculated in Excel. Statistical analyses were performed using a two-sided t-test with equal variance for all data. A p value < 0.05 is considered statistically significant.

## Acknowledgment

The authors thank all members of the Qi lab for facilitating experiments and useful discussions. The project was supported by a contract grant from Bill & Melinda Gates Foundation (INV-035661, L.S.Q). Dr. Lei S. Qi (L.S.Q.) is a Chan Zuckerberg Biohub Investigator.

## Author contributions

L.Z. and L.S.Q. conceived of the idea. L.Z. and L.S.Q. planned the experiments. L.Z. designed and cloned all the constructs and performed all the experiments and analyzed the experimental data. L.Z. and L.S.Q. wrote the manuscript. All authors read and commented on the manuscript.

## Supplementary Figures

**Supplementary Figure 1.**
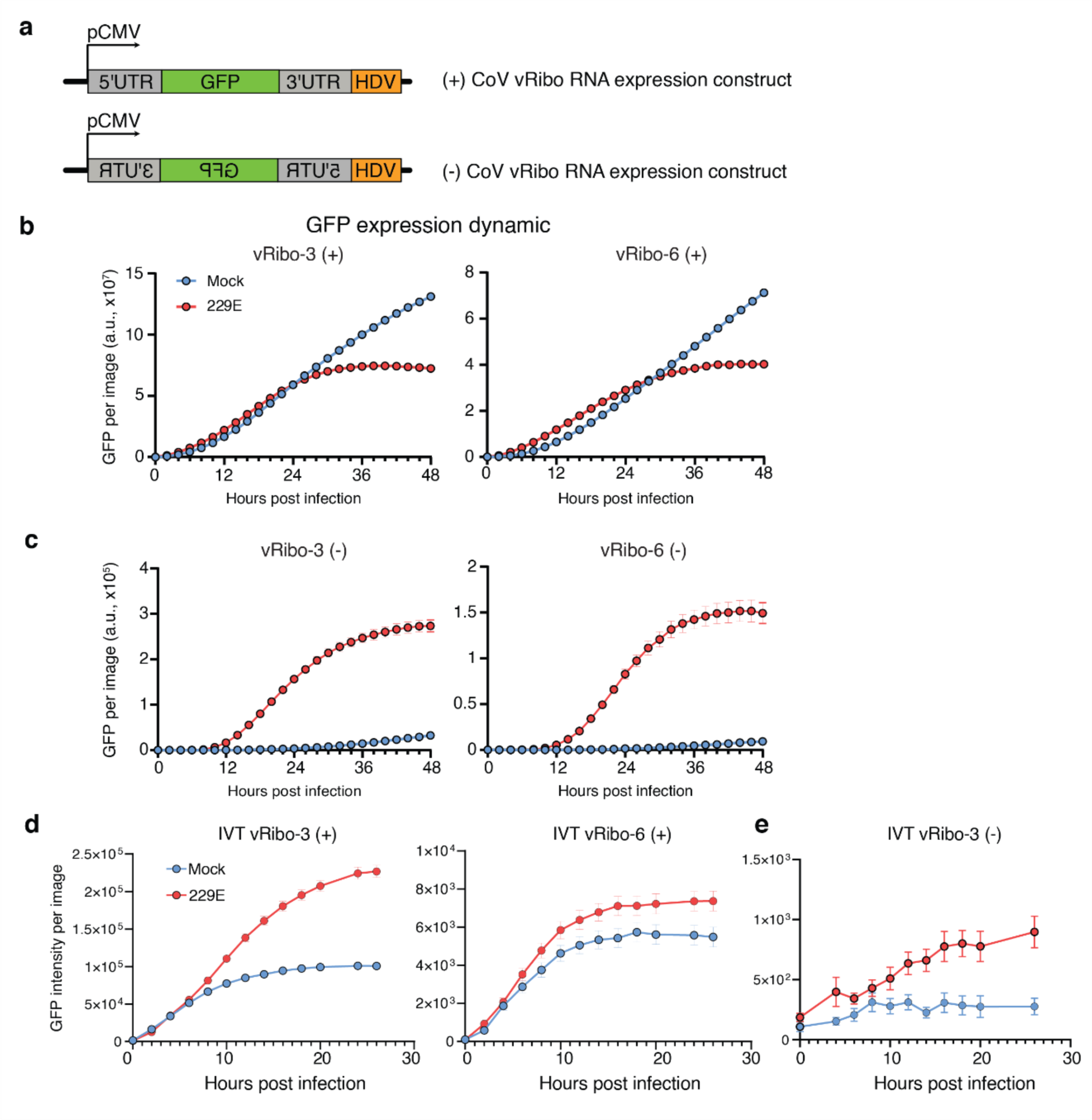
vRibos can be transcribed and replicated by human coronavirus 229E. **(a)** The construct design for expression of vRibo RNAs using plasmid DNA. **(b-e)** The GFP expression dynamics after 229E infection when the cells are transfected with plasmid DNA expressing the positive-strand (**b**) or negative-strand vRibos (**c**), and when the cells are transfected with in vitro transcribed positive-strand (**d**) or negative-strand vRibo-3 (**e**). A total of 32 images, divided into 4 separate biological replicates, were collected at each time point and the integrated fluorescence intensity was calculated.

**Supplementary Figure 2.**
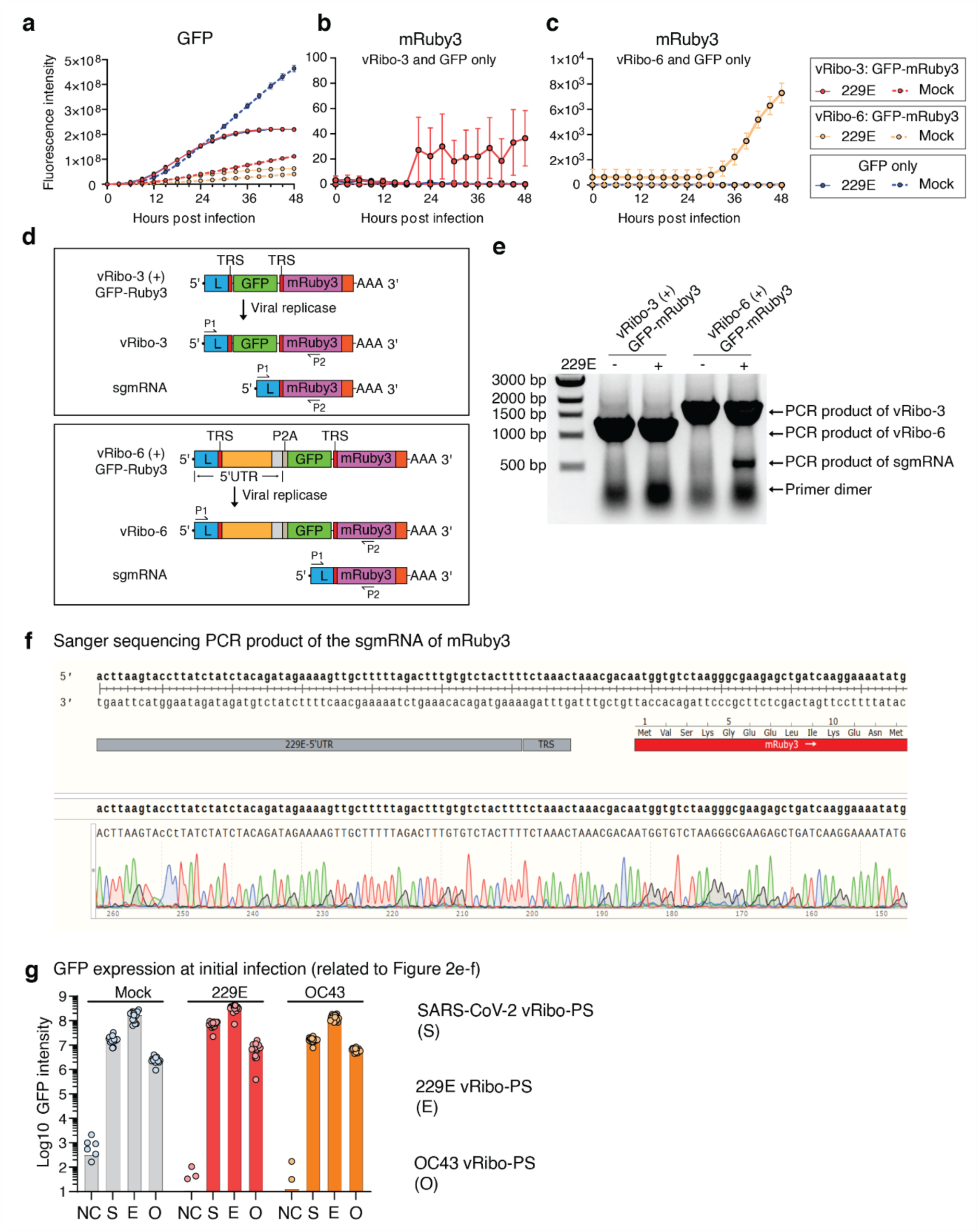
vRibos can encode a secondary protein and be packaged into progeny virions. **(a-c)** The GFP and mRuby3 expression dynamics of the 293T/hAPN cells which were transfected with the vRibos and infected with or without 229E. A total of 32 images, divided into 4 separate biological replicates, were collected at each time point and the integrated GFP and RFP fluorescence intensity was calculated separately. Data are presented as mean ± s.e.m. **(d)** The method for detection of the subgenomic transcript (sgmRNA) from vRibos using RT-PCR. **(e)** The gel electrophoresis of the RT-PCR products. **(f)** The Sanger sequencing result of the RT-PCR product of the sgmRNA of mRuby3. **(g)** The GFP expression level of the cells transfected with indicated vRibos after the initial infection. 4 independent biological replicates were performed with 4 images collected per biological replicate.

**Supplementary Figure 3.**
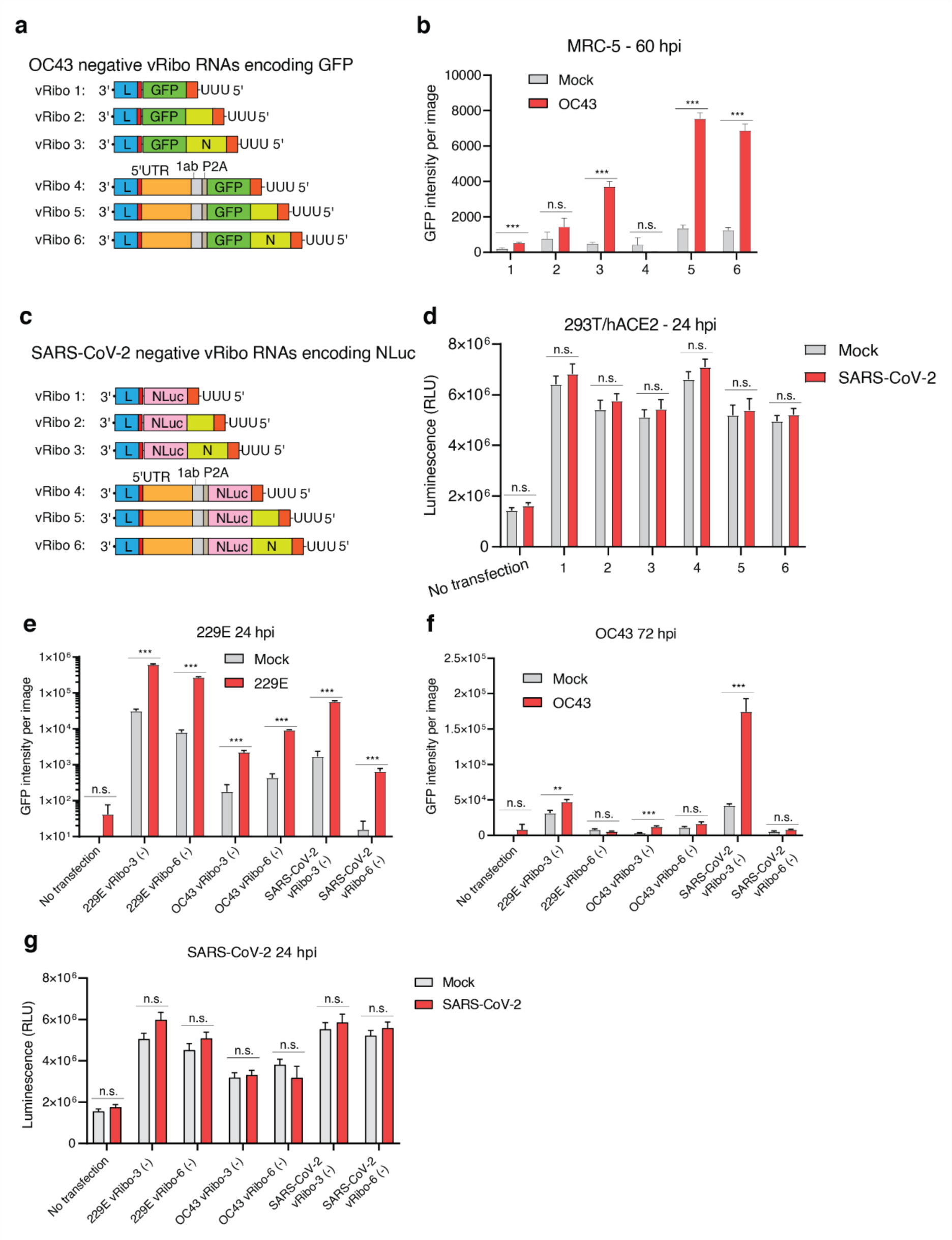
vRibos can detect and respond to broad coronaviruses. **(a)** The vRibo designs of human coronavirus OC43 and **(b)** test of the activation of the reporter expression with OC43 infection. A total of 32 images, divided into 4 separate biological replicates, were collected and the integrated GFP fluorescence intensity was calculated. **(c)** The design of the RNA sensors of SARS-CoV-2 and **(d)** test of their effectiveness with SARS-CoV-2 infection. The luciferase activity was measured with 3 independent biological replicates. **(e-g)** The reporter activation activity of vRibos of 229E, OC43, and SARS-CoV-2 by 229E **(e),** OC43 **(f)**, and SARS-CoV-2 infection. 4 independent biological replicates were performed with 2 images collected per biological replicate (e-f). The luciferase activity was measured with 3 independent biological replicates (g). Data are presented as mean ± s.e.m. P values were calculated by two-tailed Student’s t-tests. n.s., not significant; **P < 0.01; ***P < 0.001.

**Supplementary Figure 4.**
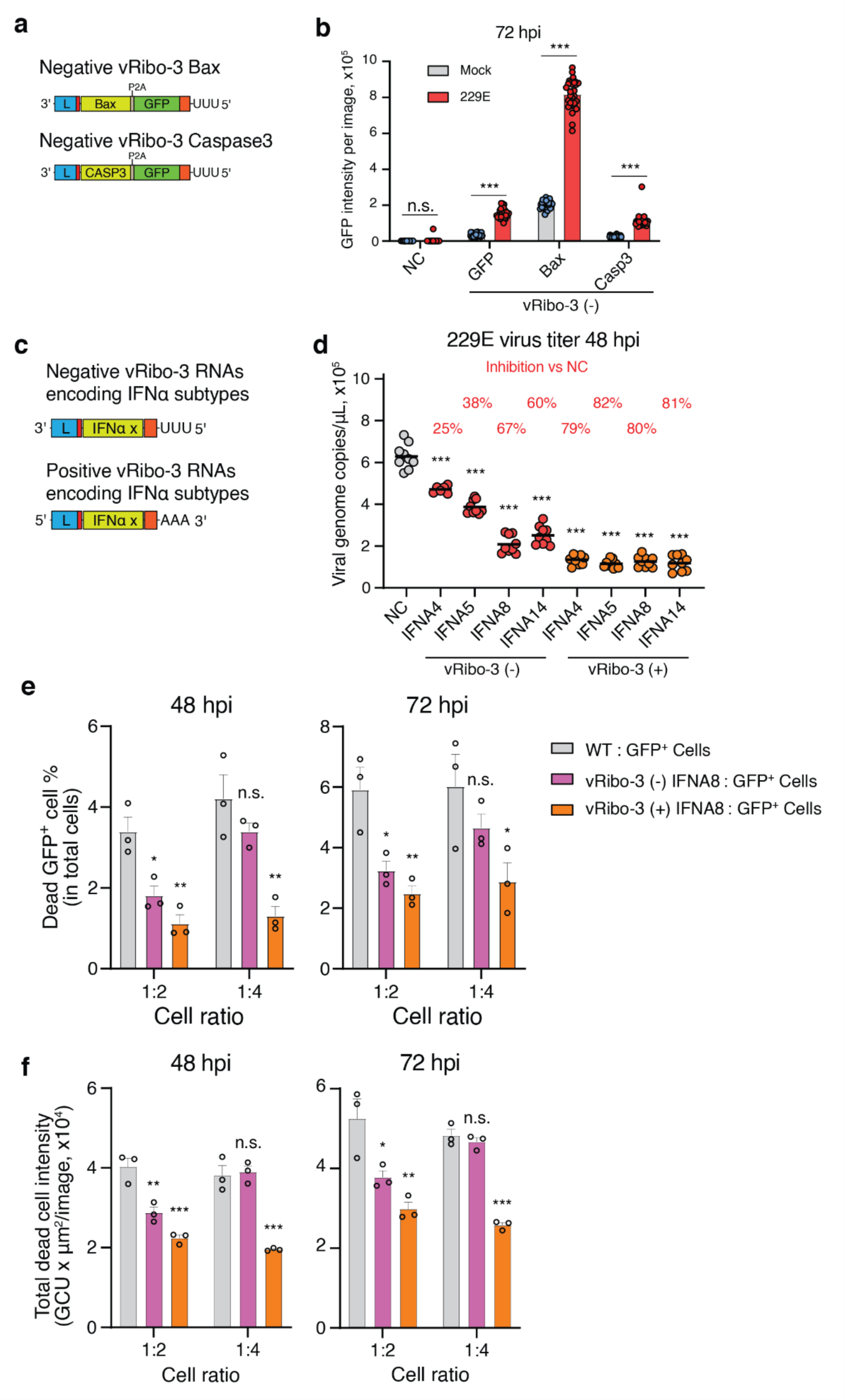
Application of vRibos for coronavirus infection detection and antiviral treatment. **(a)** The design of 229E vRibos encoding the apoptosis inducers, Bax and Caspase 3, and **(b)** the reporter GFP expression from these vRibos in mock or infected cells. 4 independent biological replicates were performed with 8 images per biological replicate. **(c)** The design of vRibos encoding the IFNs and **(d)** reduction of the virus titer at 48 hpi. 3 independent biological replicates were performed with 3 technical repeats per biological replicate. The bar represents the average of the group, while each circle represents an individual image. P values were calculated by two-tailed Student’s t-tests. **(e-f)** The percentage of dead GFP^+^ cells in total cells **(e)** and the total dead cell intensity **(f)** were measured at 48 and 72 hpi after the indicated mixtures of cells were challenged with 229E. 3 independent biological replicates were performed. Data are presented as mean ± s.e.m. P values were calculated by one-tailed Student’s t-tests. n.s., not significant; *P < 0.05; **P < 0.01; ***P < 0.001.

**Supplementary Figure 5.**
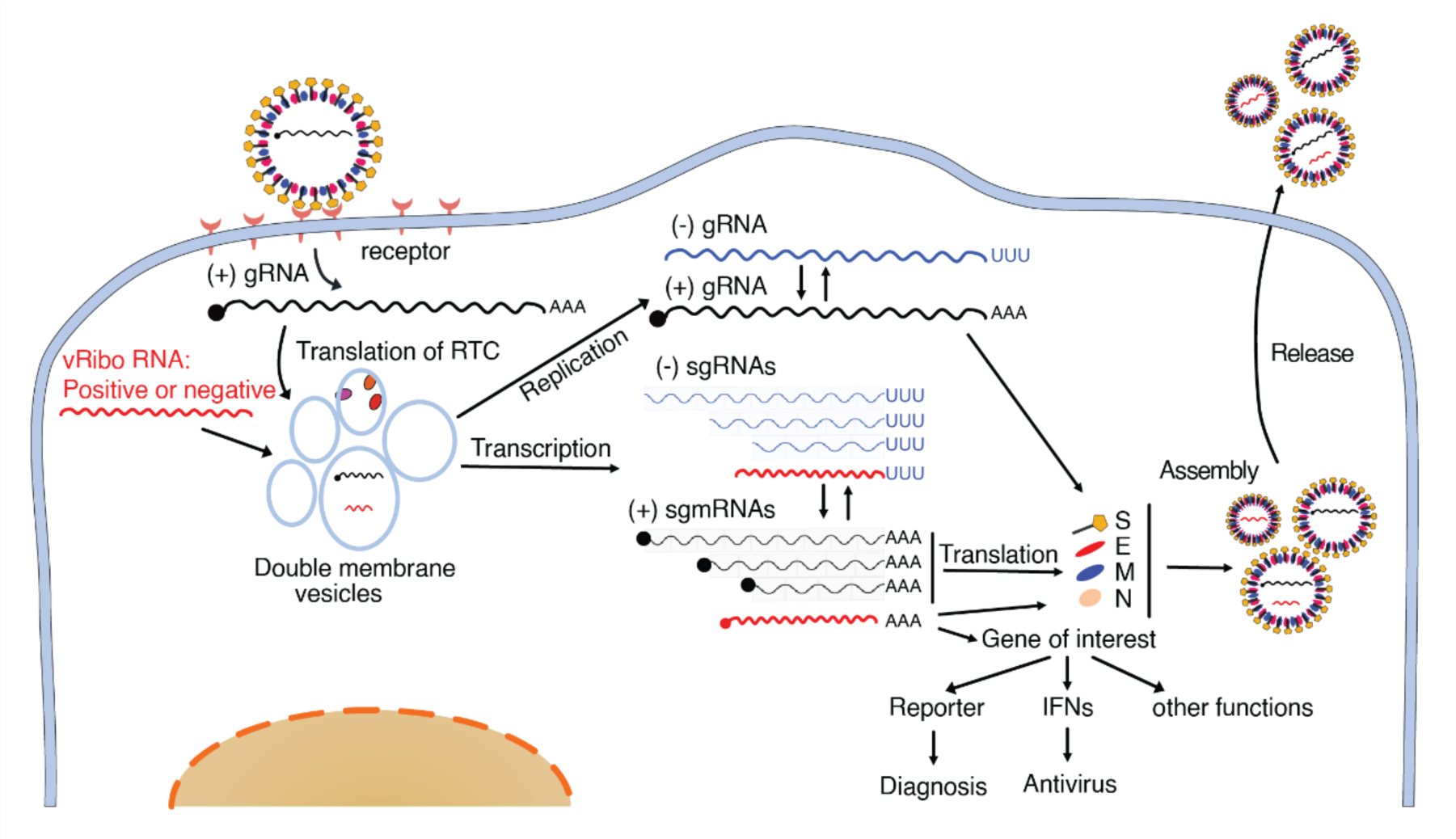
The vRibo system offers a concept by harnessing viral regulatory machineries to detect virus infection and trigger antiviral therapy.

